# Estimation of the curved body length of tuna larvae from microscope images using a zero-shot model and image processing techniques

**DOI:** 10.64898/2026.05.26.727561

**Authors:** Yuka Iwahara, Hiroshige Tanaka, Taiki Ishihara, Atsushi Tawa, Ifue Fukuchi, Masahiro Manano, Tomoya Nishino, Hiroki Yaemori, Yasutoki Shibata

**Affiliations:** Fisheries Resources Institute, National Research and Development Agency, Japan Fisheries Research and Education Agency, Yokohama, Kanagawa, Japan; Okinawa Churaumi Aquarium, Motobu, Okinawa, Japan; Computermind Corp., Nishi-Shinjyuku, Tokyo, Japan

## Abstract

Fixed larval specimens often shrink and curve, making length measurement labor-intensive. Although recent studies have demonstrated efficient fish-length estimation from images using deep learning, methods for estimating curved length remain limited. Furthermore, although deep learning is a powerful method for object detection in images, an essential step for length measurement, it requires preparing large amounts of training data, which can hinder practical implementation. In this study, we used a zero-shot model that requires no training to detect fish in an image. The curved length was then estimated using an image-processing approach that combines image thinning with Bézier curve approximation, and its accuracy was evaluated. We analyzed 1,040 larvae from five tuna species captured in stereomicroscope images. Manual measurements (notochord length or standard length; 1.5–8.5 mm) were conducted by two measurers and served as reference values. Fish regions were detected using GroundedSAM, and curved body centerlines were extracted through image thinning and approximated with Bézier curves. The curve length was used as the estimated body length. Estimation accuracy was assessed using bias and standard deviation between estimated and manual measurements. GroundedSAM detected all 1,040 fish, although there were 49 overdetections. Overdetection was caused by the double-detection of a single fish or by the misidentification of debris and light reflections as fish. Although the standard deviation of the differences between manual measurements and the image analysis–based (IAB) method was larger than the inter-measurer differences, the bias for ≤5 mm was comparable to or smaller than the inter-measurer bias. According to the strength-of-agreement criteria for the concordance correlation coefficient, the IAB method demonstrated substantial agreement in the ≤5.0-mm range. The IAB method accurately measured most curved tuna larvae without prior training, particularly in the ≤5.0-mm range. Combining the IAB method with manual remeasuring can improve the efficiency of curved-length measurement tasks.

## 1. Introduction

Recent remarkable progress in image-based deep learning techniques has spurred studies to improve measurement efficiency and reduce manual labor across various scientific fields (Masud et al., 2020; Ni et al., 2026; Wang et al., 2023). In fisheries science, this approach enables efficient collection of biological data, including species, body length, weight, and age, and has therefore been used in studies on monitoring, stock assessment, and stock management (Lekunberri et al., 2022; Moen et al., 2018; Palmer et al., 2022; Shibata et al., 2024).

Fish length data are among the most fundamental biological parameters in fisheries science (Bunch et al., 2013; Miranda et al., 2024). In stock assessment, length data are required to infer a population’s age structure through a length–age relationship (Piner et al., 2016). Length can also be converted to body weight, which is essential information for aquaculture. (Muñoz-Benavent et al., 2018). In larval studies, identifying growth stages based on length helps elucidate fundamental ecological processes (Tanaka et al., 2006). Therefore, numerous studies have used deep learning to efficiently extract critical length information from images. For instance, Shibata et al. (2024) estimated total length by segmenting the fish body, generating a minimum bounding box for each segment and using the longest side of the box as the total length. Tanhir and Iqbal (2024) estimated fork length using key point detection. Álvarez-Ellacuría et al. (2020) estimated total length by detecting only the head region and converting head length to total length. These approaches generally assume that the fish body or the measurement region is straight.

However, fish are not always photographed in a straight posture; some are captured in a curved posture. This can occur in elongated-bodied species, swimming fish, and fixed larvae (Esselman et al., 2025; Ishihara et al., 2025; Muñoz-Benavent et al., 2018; Takizawa et al., 1994). Previous studies have used image processing and geometric models to estimate the curved length of swimming fish (Garcia et al., 2020; Muñoz-Benavent et al., 2018; Zhou et al., 2023). Muñoz-Benavent et al. (2018) developed a specialized geometric model to estimate the snout-to-fork length of Atlantic bluefin tuna (*Thunnus thynnus*) from an overhead perspective, achieving a measurement error of ±3% for 90% of the fish. Another approach focused on swimming fish photographed from the side (Garcia et al., 2020). In this method, image thinning, an image-processing technique, was applied to skeletonize a fish mask, and the resulting skeletonized line was smoothed using the RANSAC algorithm to obtain a curve corresponding to curved body length. This approach enables species-independent estimation of curved body length.

However, liquid-immersed larvae fixed with agents such as ethanol or formalin can exhibit even more complex body curvatures than those observed during swimming (Ishihara et al., 2025; Takizawa et al., 1994). Conventionally, this measurement has been performed manually using software such as ImageJ (Ishihara et al., 2025). This process requires drawing a curve for each larva, making it extremely labor-intensive for large sample sizes. Furthermore, manual measurements introduce the potential for inter-measurer variability. Therefore, an automated method is necessary for estimating the length of curved bodies, including ethanol-dehydrated larvae. An effective approach for approximating more complex shapes is to use Bézier curves (Bézier, 2014). They are commonly employed in computer graphics and related areas, such as computer-aided geometric design (Baydas and Karakas, 2019; Elradi et al., 2024), and even presentation software, such as Microsoft PowerPoint, uses them for drawing smooth, editable shapes (https://learn.microsoft.com/en-us/office/vba/api/powerpoint.shapes.addcurve). Applying such curve-fitting techniques may enable the estimation of body length in ethanol-dehydrated larvae.

To estimate fish length from images, the first step is to detect the fish region. Rasmussen et al. (2022) used image-processing techniques to detect the fish region. However, they noted that detection accuracy was affected by image quality and suggested that deep learning should be introduced to mitigate this. Although deep learning is a powerful method for object detection, it generally requires a large amount of training data, which can hinder its practical implementation. To address this limitation, techniques such as few-shot and one-shot learning, which require far fewer training images, have been developed in recent years (Hsieh et al., 2019; Zhang et al., 2023). In addition, foundation models, which undergo extensive pretraining, have recently emerged (Bommasani, 2021). These models enable zero-shot detection and segmentation, allowing various objects to be detected and segmented without training on a custom dataset (Ren et al., 2024). GroundedSAM (Ren et al., 2024) achieves zero-shot segmentation by combining object detection from GroundingDINO (Liu et al., 2024) with segmentation from the Segment Anything Model (SAM) (Kirillov et al., 2023). For example, the model can detect fish regions without prior training by simply being given a text prompt such as “fish.” Hasegawa and Nakano (2024) reported that GroundedSAM could detect adult fish and noted that accurate masking was possible when fish did not overlap. This capability may facilitate the introduction of region detection for larvae on glass slides. However, no attempt has been made to detect fish larvae.

Our aim is to automate the measurement of body length in fixed curved larvae. To achieve this, we first detect and segment the fish region using a zero-shot model and then evaluate detection performance and mask accuracy. Subsequently, we estimate the length of curved fish using image processing techniques based on image thinning and Bézier curve approximation and finally evaluate the performance of the body length estimation.

## 2. Materials and methods

### 2.1 Fish selection and image acquisition

This study focused on five species of tuna larvae (family Scombridae, tribe Thunnini): Pacific bluefin tuna *Thunnus orientalis*, yellowfin tuna *Thunnus albacares*, skipjack tuna *Katsuwonus pelamis*, bullet tuna *Auxis rochei*, and frigate tuna *Auxis thazard*. The larvae were collected using ring nets (diameter, 2 m; mesh size, 0.33 or 1.00 mm) in Japanese coastal waters from 2017 to 2024 and fixed onboard in 99.5% ethanol. Larval images were captured at 0.8–5.6× magnification using a digital camera (DP23-AOU, Olympus) mounted on a stereomicroscope. Fish were placed individually or in groups on graduated or grid-marked glass slides, avoiding overlap (Supplementary Figure 1). We used body-length classes (0.5-mm bins) that contained 20 or more individuals for each species. From each class, 20 individuals were randomly selected to avoid bias across collection years, resulting in 1,040 individuals evaluated (Supplementary Table 1). The body length used for this selection process was measured in advance using the same method as in Section 2.2. Fish with partially collapsed bodies, looped bodies, or three-dimensional bends were excluded from the analysis, as they were outside the scope of this study.

### 2.2 Manual measurement

To obtain manual length measurements for accuracy evaluation, body length (either notochord length or standard length) was measured using ImageJ (Abràmoff et al., 2004). Grid or scale markings printed on the glass slide and visible in digital images were used for scaling. Two people measured the same fish to evaluate the accuracy of our image analysis. Both measurers were highly skilled, with >3 years of experience. Although the notochord length and standard length refer to the same anatomical region, the name of that region changes with the growth stage (Shimizu and Shiozawa, 2004) (Supplementary Figure 2).

### 2.3 Detection and segmentation of the fish region

Fish regions were detected and segmented using GroundedSAM (Ren et al., 2024). GroundedSAM combines object detection with GroundingDINO (Liu et al., 2024) and segmentation with SAM (Kirillov et al., 2023).

For GroundingDINO (version 0.1.0), the text prompt was set to “fish.” The box threshold was varied in 0.05 increments from 0.30 to 0.95 to select a value that would prevent underdetection. All other parameters were left at their default settings. We used the pretrained GroundingDINO swinb cogcoor checkpoint. For SAM (version 1.0), the bounding boxes selected above were used as prompts. All other parameters were left at their default settings. We used the pretrained checkpoint sam_vit_h_4b8939. The development environment consisted of Python 3.9.16 and PyTorch 2.2.0 + cu121. The GPU used was an NVIDIA Quadro RTX 8000.

To evaluate fish detection accuracy, we counted both overdetection and underdetection. We used intersection over union (IoU) to determine the success of detection (Tang and Yuan, 2015):

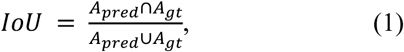

where *A_pred_* and *A_gt_* denote areas in the predicted and ground-truth masks, respectively. If IoU exceeded 0.5, the detection was marked “correct.” The correct masks for IoU calculation were obtained by manually refining the automatically generated masks in Method 2.3. The refinement was performed using the FIAS-Deep Labeler annotation tool (version 1.3.3, Computermind Corp.), which was developed to facilitate easy annotation of fish images. In addition, we calculated the IoU for each fish to evaluate the accuracy of segmentation mask. In this process, we excluded overdetected masks. In this study, “mask” refers to a binary mask representing the fish region, obtained either from a segmentation model or created manually.

### 2.4 Extraction of the curve along the curved fish body

In our image analysis–based (IAB) measurement, the centerline of the curved fish body was obtained by skeletonizing the mask using image thinning and then approximated by a Bézier curve. The length of the resulting curve was calculated and used as the estimated body length.

Image thinning was applied to the fish mask (Figure 1a), followed by pruning to retain only the longest skeletonized line (Figure 1b). A Bézier curve was then fitted to this pruned curve (Figure 1c). The number of control points was set to the number of points in the pruned curve divided by 20. However, our preliminary analysis showed that in small fish (≤4.5 mm) with underdeveloped skeletons, more complex bends can occur over short distances. Therefore, for such cases, the number of control points was increased to the number of points in the pruned curve divided by 5. In both settings, when the number of points in the pruned curve was fewer than 40 or 10, respectively, the number of control points was set to half the number of points in the pruned curve.

**Figure 1.**
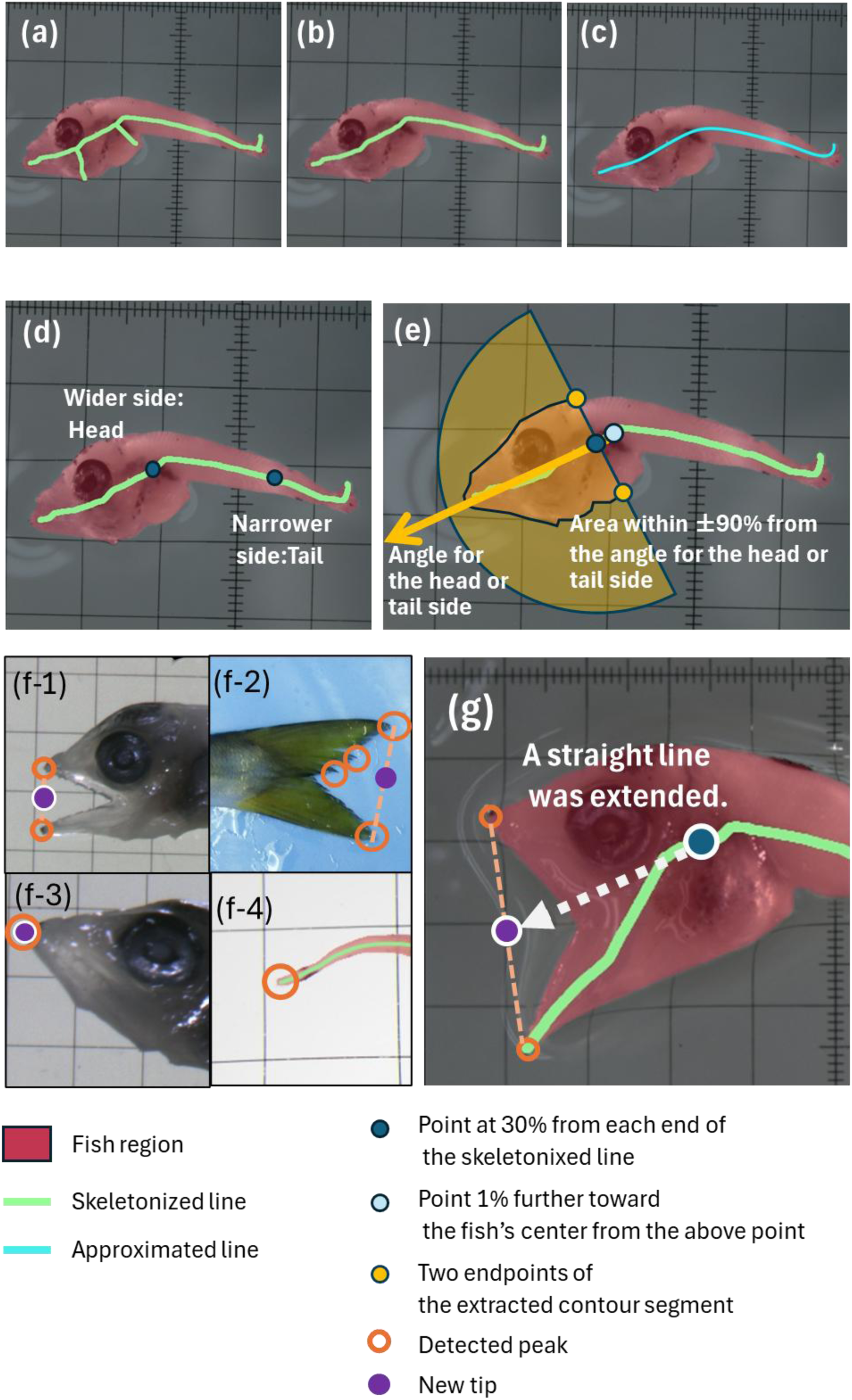
Images illustrating the process for estimating fish body length. (a) Curve obtained by image thinning. (b) Curve connecting the two longest points selected by pruning. (c) Curve obtained by Bézier curve approximation. (d) Definition of the head and tail: the wider side is defined as the head, and the narrower side as the tail. (e) Angle toward the head or tail side and the region within ±90° used to determine mask width. (f-1 and f-2) Start and end point positions when multiple peaks are detected; the centers of the farthest peaks at each end are used. (f-3 and f-4) Case in which a single peak is detected. For the head side (f-3), the peak is used as the starting point; for the tail side (f-4), no new endpoint is defined, and the point from the original approximation line is used. (g) Case in which new tups are defined. A straight line is extended from the 30% position of the approximation line to the new tip on the head side, and from the 5% position to the new tip on the tail side. (f-2) is shown using an adult fish image. Because the IAB method is a general-purpose length estimation technique that can be applied from larvae to adults with only minor parameter adjustments, this step is rarely necessary for standard body lengths of larvae but is frequently used for total length estimation in adult fish.

However, the above processes sometimes resulted in two issues that affected the accuracy of length estimation: (1) the approximation line did not extend the mask boundaries, which could occur at both ends and (2) the approximation line on the caudal fin exhibited unnecessary bending (Figure 2). Therefore, the following additional processing was applied before the approximation. Because the processing differs between the head and tail masks, as explained below, we first determined which side of the mask corresponded to the head or tail. We measured the width of the mask at points 30% from each end of the longest skeletonized line and defined the wider side as the head and the narrower side as the tail (Figure 1d). To calculate the mask width, we first determined the angle for the head or tail side by connecting two points: one at the 30% position and another 1% further toward the fish’s center (Figure 1e). Then, we extracted the portion of the mask contour whose direction fell within ±90° of this angle (Figure 1e). Finally, we calculated the width by measuring the distance between the two endpoints of the extracted contour segment (Figure 1e). Peaks were then detected using the masks at both ends. If multiple peaks were detected, the average position of the two farthest peaks was used as the tip (Figures 1f1 and 1f2). If only one peak was detected, the original longest skeletonized line was used directly for the tail (Figure 1f4), and the detected peak was used as the tip for the head (Figure 1f4). In rare cases where no peak could be detected, the original longest skeletonized line was used. When a new tip was defined, a straight line was extended from the point located 30% along the longest skeletonized line for the head and from the 5% point for the tail (Figure 1g). After this processing, the resulting line was approximated with a Bézier curve. Finally, the length of the resulting curve in pixels was measured. Body length was calculated by multiplying this pixel length by the length-per-pixel value (mm/pixel), which was measured separately.

**Figure 2.**
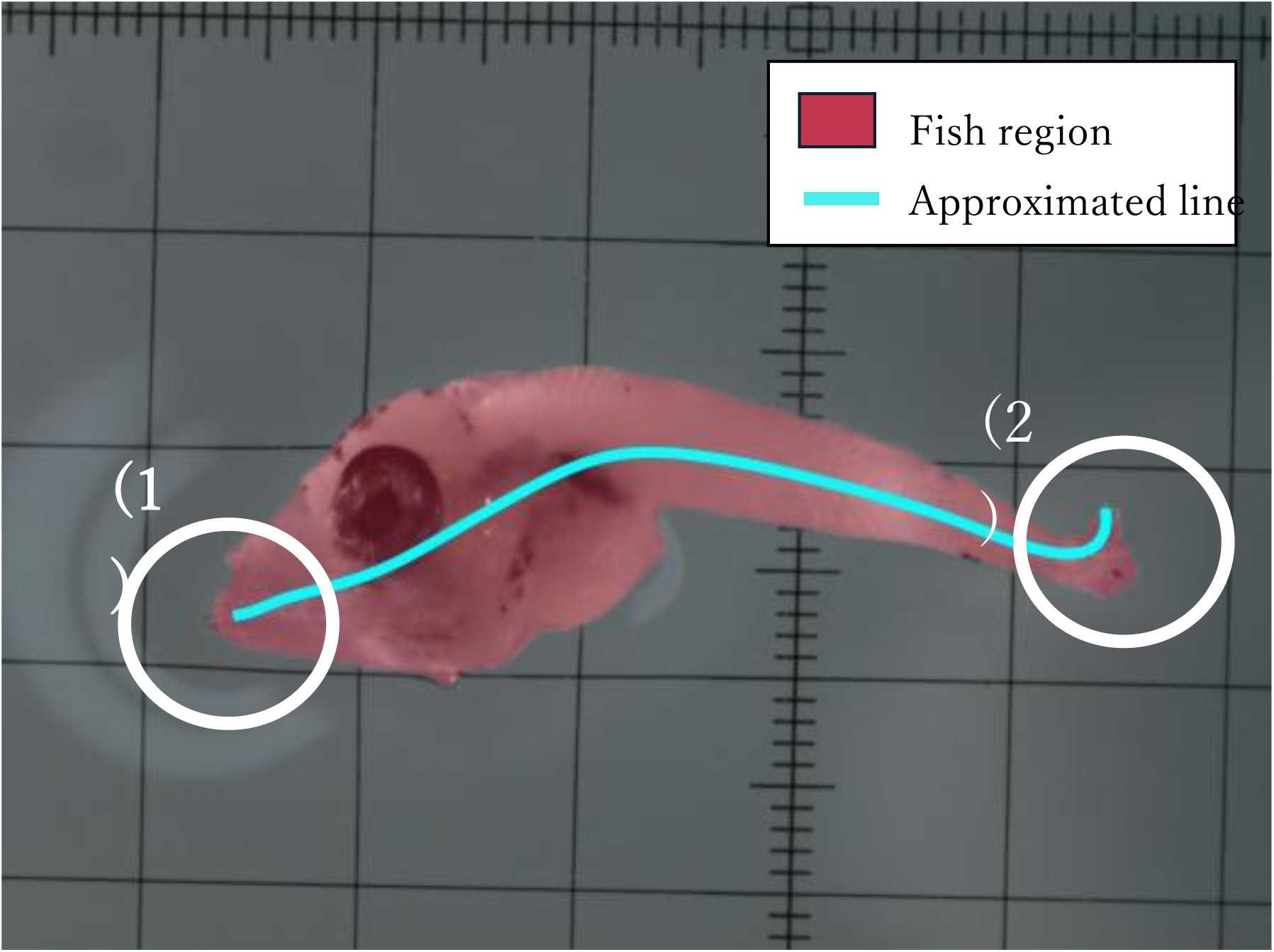
Two issues remained after the approximation process with a Bézier curve: (1) the approximation line did not extend to the mask boundaries, which could occur at both ends and (2) the approximation line on the caudal fin exhibited unnecessary bending.

However, in manual measurements, the measurement typically starts at the tip of the upper jaw. To assess the impact of this difference, we also performed an accuracy evaluation using IAB measurements that started at the tip of the upper jaw. For this evaluation, the peak corresponding to the upper jaw among the two detected peaks was used as the starting point for the head. However, because the fish were not consistently oriented and we had no automated way to identify which peak corresponded to the upper jaw, we first manually confirmed whether the upper jaw corresponded to the side with the larger or smaller y-axis value and selected the corresponding side. In actual operation, we assume that the fish will be positioned with the upper jaw at the top and that the side with the larger y-axis value (Supplementary Figure 3a) will therefore always be selected, allowing the upper jaw to be automatically selected.

For fully automated measurement in the future, scale detection must be automated as well. However, due to the limited sample size, previously photographed images were used in this study, and the grid or scale markings printed on the glass slides in these images were not well suited for automated image processing. Therefore, the scales were manually identified by a person other than measurers A and B using FIAS-Deep Labeler annotation tool. Two points where the grid or scale markings were clearly visible were identified, and the pixel distance between these points was divided by the actual length (mm) to obtain the length-per-pixel value (mm/pixel).

### 2.5 Calculation of the difference in measurement values between manual and IAB measurements

To evaluate the accuracy of the IAB measurements, we calculated the relative and absolute biases, as well as the standard deviation of the measurement differences between the manual and IAB methods. In addition, some studies use measurement differences between measurers as an index of acceptable differences in AI measurements (Moen et al., 2018). Therefore, we also calculated the relative and absolute biases, as well as the standard deviation of the measurement differences between two measurers, and compared these values with those of the IAB method. First, relative and absolute differences were calculated as follows:

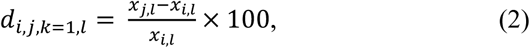

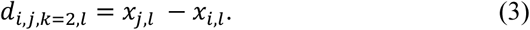

*d_k=_*_1_ and *d_k=_*_2_ denote the relative and absolute differences, respectively. *x* denotes the length measurement value, with *i* representing each measurement source (𝑖 = 1 for measurer A, 𝑖 = 2 for measurer B, and 𝑖 = 3 for the IAB method) and *l* representing each fish (*l* = 1, …, 1,040). The relative and absolute differences were calculated for each pair of indices (𝑖, 𝑗), where 𝑖, 𝑗 ∈ {1,2,3} and 𝑖 ≠ 𝑗, corresponding to three comparisons: measurer A vs. measurer B, measurer A vs. the IAB method, and measurer B vs. the IAB method. The bias (*b*) and standard deviations (*s*) for each relative and absolute difference were calculated as follows:

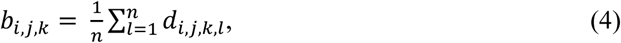

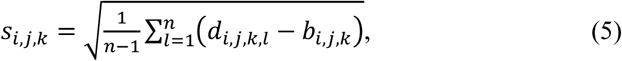

where *n* denotes the total number of fish (*n* = 1,040). Section 3 primarily presents absolute differences, with relative differences provided as supplementary information.

To understand the relationship between measurement differences and body length, the data were visualized using a scatter plot. To identify trends in this relationship, segmented regression analysis was performed to detect breakpoints at which the relationship changed. The significance of these change points was assessed using the Davies test. These analyses were conducted in R version 4.4.0 (R Core Team, 2024) using the “segmented” package.

To assess interspecific differences in measurement differences between measurers and the IAB method, we used the Kruskal–Wallis test (Kruskal and Wallis, 1952). Before conducting these two tests, normality was assessed using the Shapiro–Wilk test (Shapiro and Wilk, 1965), which indicated that the data were non-normal and supported the use of these statistical methods (Supplementary Table 2). This test was conducted within the range common to all species (3.0–6.5 mm), because the maximum and minimum bin widths varied among species (Supplementary Table 1).

### 2.6 Assessment of agreement

We used the concordance correlation coefficient (CCC) as an evaluation index. The CCC is one of the most common approaches used for assessing agreement among observers or instruments (Carrasco et al., 2013). It has been used to compare manual and AI measurements(Rizzo et al., 2017; Wieland and Heuwieser, 2025).

The CCC was calculated as follows (Lawrence and Lin, 1989):

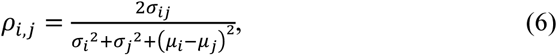

where *σ_ij_* denotes the covariance, 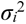 and 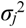 denote the variances, and *μ_i_* and *μ_j_* denote the means. We introduced the same index 𝑖 in Section 2.5 to denote each measurement source: 𝑖 = 1 for measurer A, 𝑖 = 2 for measurer B, and 𝑖 = 3 for the IAB method. The CCC was calculated for each pair of indices *i* and *j*, where 𝑖, 𝑗 ∈ {1,2,3} and 𝑖 ≠ 𝑗, corresponding to the three comparisons: measurer A vs. measurer B, measurer A vs. the IAB method, and measurer B vs. the IAB method. McBride (2005) categorized the strength-of-agreement criteria for the CCC based on their simulation as follows: almost perfect (>0.99), substantial (0.95–0.99), moderate (0.90–0.95), and poor (<0.90). These analyses were performed in R using the “epiR” package. The CCC calculation was performed separately for the ranges before and after the breakpoint, if a breakpoint was detected using the segmented regression described in the previous section.

## 3. Results

### 3.1 Accuracy of detection number and masks

Adjusting the box prompt value changed the number of overdetection and underdetection. When the box prompt value was set to 0.3, all 1,040 fish were detected, with none undetected and 49 overdetected (Table 1). The breakdown of overdetections comprised 19 misdetections of debris or plankton, 19 double detections (Figure 3a), and 10 misdetections of water reflections (Figure 3b). IoU was 0.9 or higher for all fish except two, indicating accurate segmentation (Figure 4).

**Figure 3.**
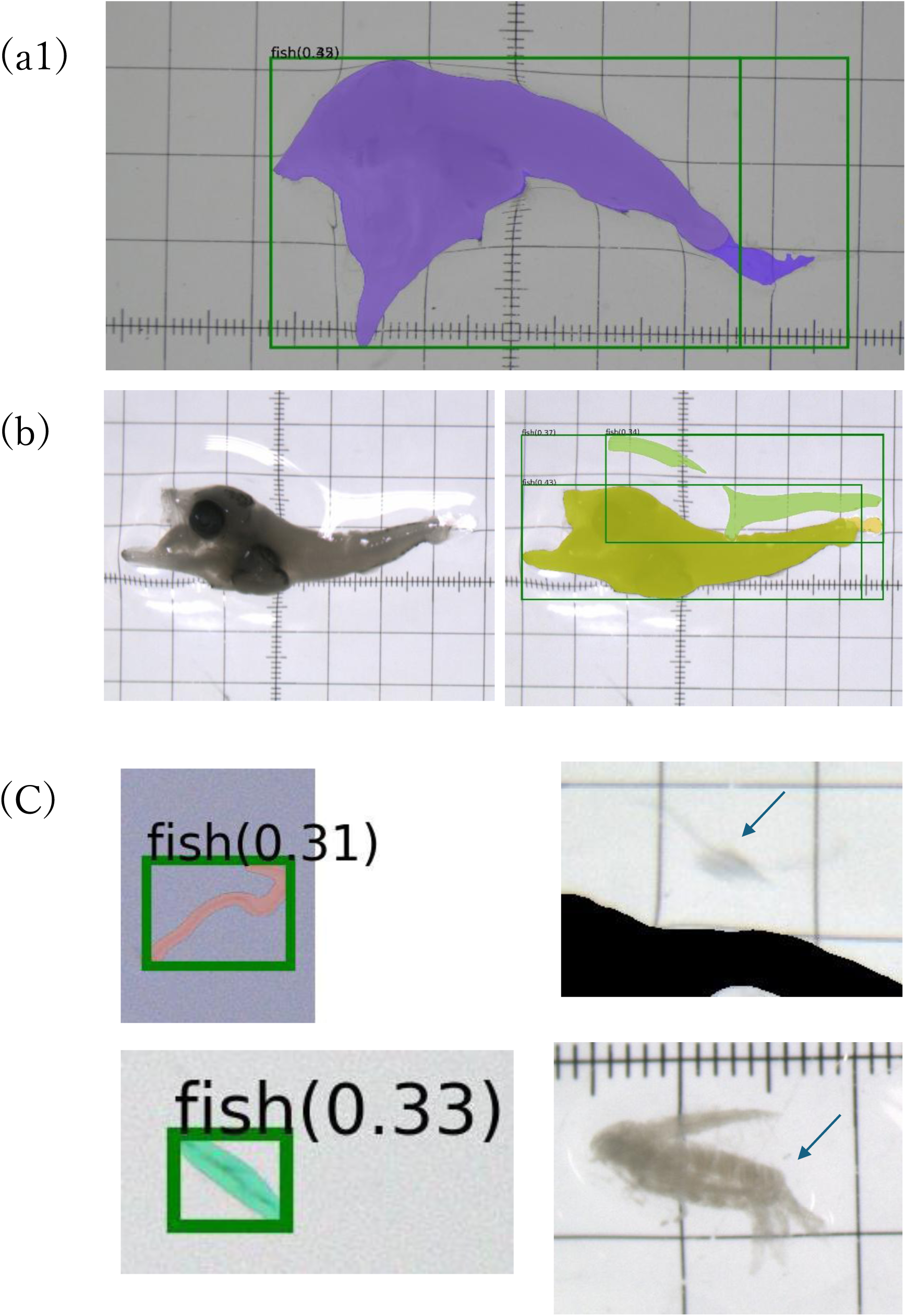
Examples of overdetections: (a) double detection in a single fish, (b) misidentification of light reflections as fish, and (c) misidentification of plankton or debris as fish.

**Figure 4.**
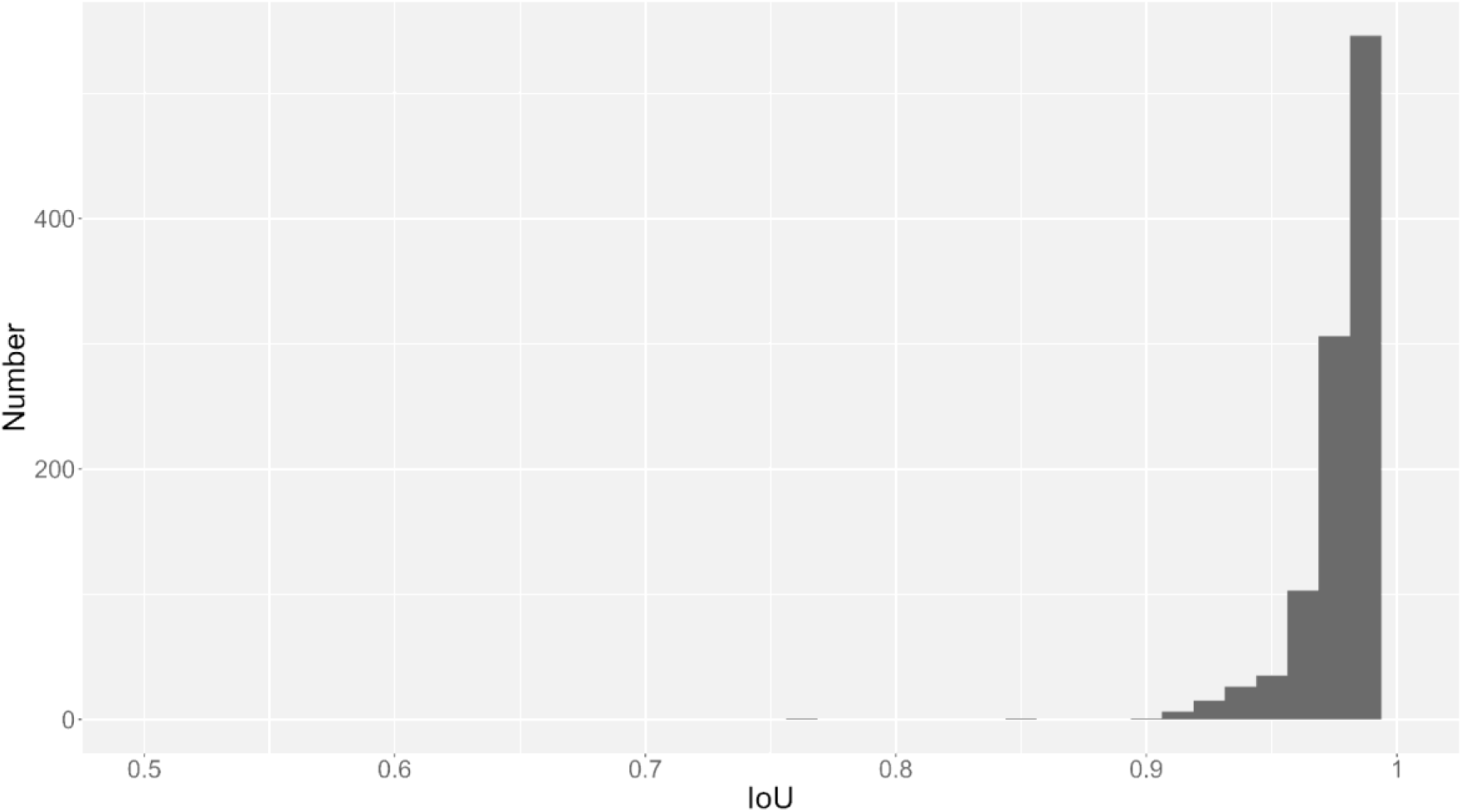
Histogram of IoU values for the detected fish masks. Only correctly detected masks were included in the calculation.

**Table 1.** Number of true detections, overdetections, and underdetections of fish when the box prompt value was changed.

| Box prompt | 0.3 | 0.35 | 0.4 | 0.45 | 0.5 | 0.55 | 0.6 | 0.65 | 0.7 | 0.75 | 0.8 | 0.85 | 0.9 | 0.95 |
| --- | --- | --- | --- | --- | --- | --- | --- | --- | --- | --- | --- | --- | --- | --- |
| True detection | 1040 | 1039 | 1034 | 1022 | 998 | 927 | 803 | 631 | 454 | 282 | 165 | 91 | 57 | 0 |
| Over detection | 49 | 30 | 9 | 2 | 1 | 0 | 0 | 0 | 0 | 0 | 0 | 0 | 0 | 0 |
| Under detection | 0 | 1 | 6 | 18 | 42 | 113 | 237 | 409 | 586 | 758 | 875 | 949 | 983 | 1040 |

### 3.2 Evaluation of measurement difference

The difference between manual measurements and the IAB method showed a bimodal pattern from around 5 mm (Figure 5, Supplementary Figure 4). One mode was centered near zero, whereas the other showed increasing positive differences with increasing body length (Figure 5). Although the maximum and minimum bin widths varied among species (Supplementary Table 1), no significant interspecific differences were observed within the range common to all species (3.0–6.5 mm) (Figure 5, Table 2). Similar tendencies were observed when the starting position was at the upper peak (Supplementary Figure 5). Segmented regression identified breakpoints of 5.00 mm (standard error 0.34) for measurer A and the IAB method, and 5.34 mm (standard error 0.32) for measurer B and the IAB method, where the slope of the relationship between body length and absolute difference changed (Table 3, Figure 6).

**Figure 5.**
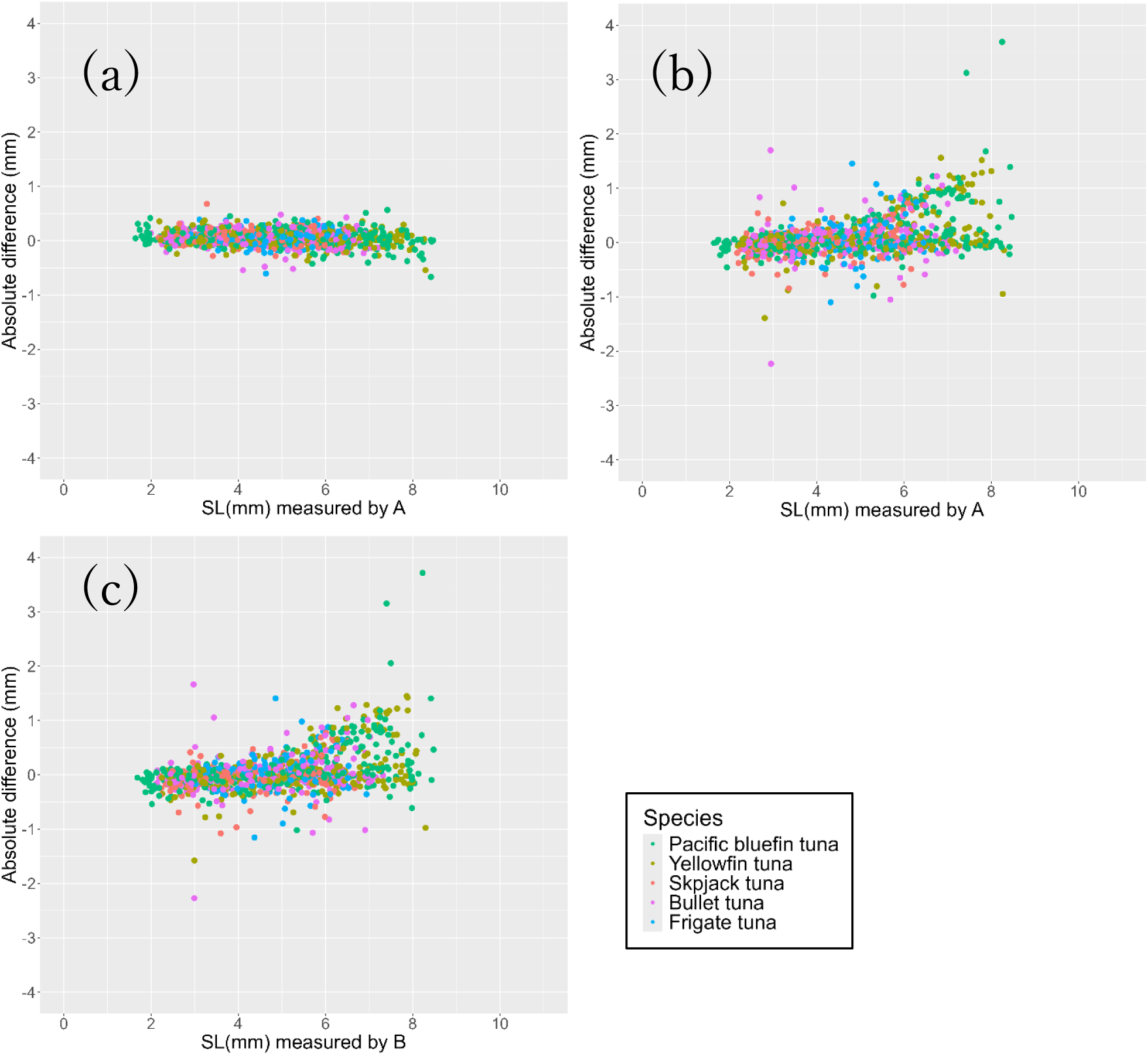
Scatter plots of body length versus absolute differences: (a) measurers A and B, (b) measurer A vs. the IAB method, and (c) measurer B vs. the IAB method. Common and scientific names of the five species: Pacific bluefin tuna *Thunnus orientalis*, yellowfin tuna *Thunnus albacares*, skipjack tuna *Katsuwonus pelamis*, bullet tuna *Auxis rochei*, and frigate tuna *Auxis thazard*.

**Figure 6.**
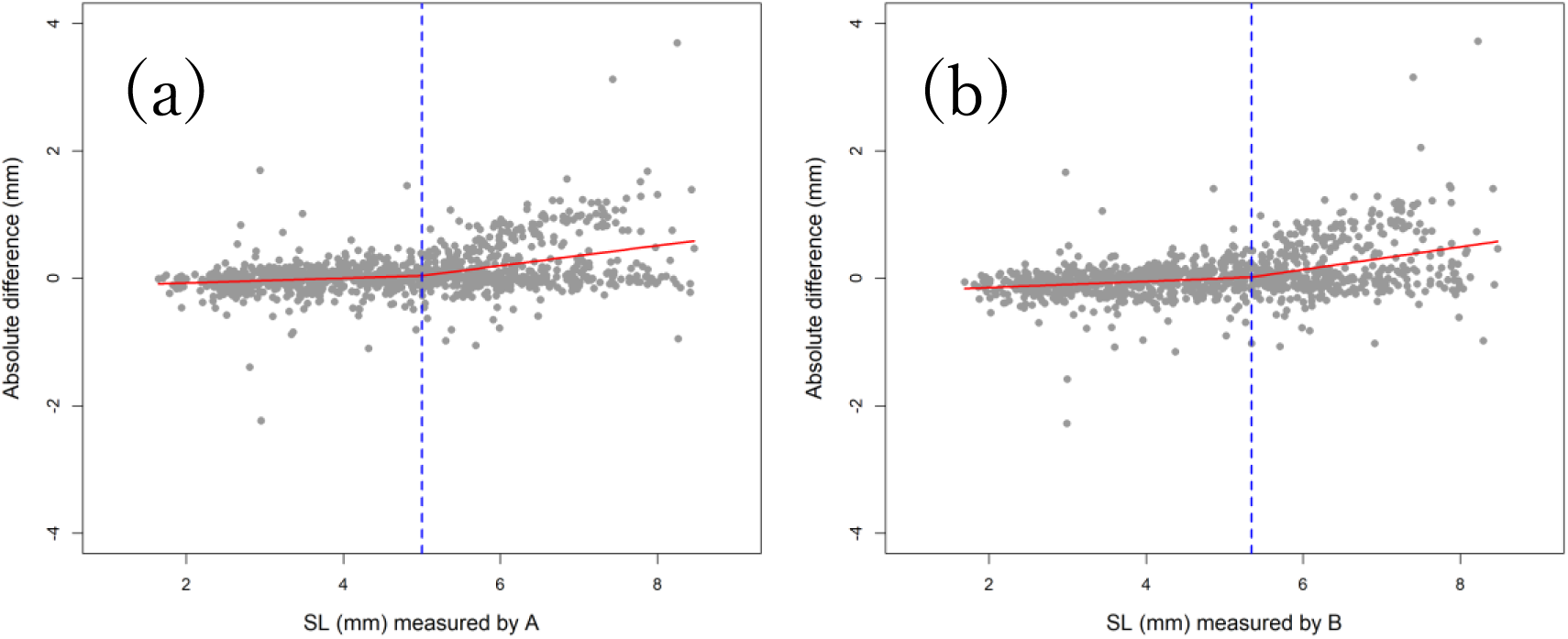
Changing point and scatter plot showing body length and measurement differences: (a) measurer A vs. the IAB method and (b) measurer B vs. the IAB method. Gray points indicate individual measurement differences, and red dashed lines indicate the fitted segmented regression. The bold blue dashed line marks the estimated breakpoint.

**Table 2.** Statistical test results for interspecific measurement differences between measurers and the IAB method. The Kruskal–Wallis test detected no significant differences in either the A–IAB or B–IAB comparisons for both relative and absolute differences.

|  | Relative bias |  | Absolute bias |  |
| --- | --- | --- | --- | --- |
|  | A vs IAB | B vs IAB | A vs IAB | B vs IAB |
| Chi-squared | 2.054 | 3.139 | 1.578 | 2.313 |
| df | 4 | 4 | 4 | 4 |
| P value | 0.726 | 0.535 | 0.813 | 0.678 |

**Table 3.** Results of segmented regression. “IAB” denotes the image analysis–based method. The two types of differences—between measurer A and IAB and between measurer B and IAB—were modeled using segmented regression. “Body length 1” denotes the slope before the break point and “Body length 2” denotes the change in slope after the break point. The slope after the break point is obtained by adding the estimate for “Body length 2” to that for “Body length 1.” Because the p-value for “Body length 2” was not computed, significance was assessed using the Davies test. \**p*-value < 0.05.

| Difference | Break-point |  | Coefficients of the linear terms: |  |  |  |  | Davies test |  |  |
| --- | --- | --- | --- | --- | --- | --- | --- | --- | --- | --- |
|  | Estimate | Standard error | Coefficient | Estimate | Standard error | t value | p value | Best | p value |  |
| A - IAB | 5.00 | 0.34 | Intercept | -0.15 | 0.06 | -2.33 | 0.02 * | 4.69 | 9.22.E-05 | * |
|  |  |  | Body length 1 | 0.04 | 0.02 | 2.16 | 0.03 * |  |  |  |
|  |  |  | Body length 2 | 0.12 | 0.03 | 4.70 | - |  |  |  |
| B - IAB | 5.34 | 0.32 | Intercept | -0.24 | 0.06 | -4.08 | 4.79.E-05 * | 5.49 | 4.92.E-05 | * |
|  |  |  | Body length 1 | 0.05 | 0.02 | 3.19 | 1.49.E-03 * |  |  |  |
|  |  |  | Body length 2 | 0.13 | 0.03 | 4.68 | - |  |  |  |

Across the two aggregation conditions (≤5 and >5 mm), the length difference between manual measurements and the IAB method was comparable to or smaller than the inter-measurer difference in terms of bias for ≤5 mm. By contrast, the standard deviation of the differences was larger in both conditions (Table 4, Supplementary Table 3). The absolute bias was 0.59 mm between measurers, whereas it was −0.014 and −0.073 mm between each measurer and the IAB method for ≤5 mm (Table 4). For >5 mm, the corresponding values were 0.038, 0.231, and 0.192 mm (Table 4). The standard deviation of the absolute differences was 0.129 mm between measurers, compared with 0.244 and 0.247 mm between measurers and the IAB method for ≤5 mm (Table 4). For >5 mm, the corresponding values were 0.156, 0.463, and 0.474 mm (Table 4).

**Table 4.** Bias and standard deviation of the differences between measurers A and B and between each measurer and the IAB method (IAB) under the three aggregation conditions (the entire length range, ≤5 mm, and >5 mm).

| Species | Body length range | Absolute bias (mm) |  |  | Standard deviation of absolute difference (mm) |  |  |
| --- | --- | --- | --- | --- | --- | --- | --- |
|  |  | A vs B | A vs IAB | B vs IAB | A vs B | A vs IAB | B vs IAB |
| All | All | 0.049 | 0.096 | 0.046 | 0.142 | 0.379 | 0.390 |
|  | 5mm<= | 0.059 | -0.014 | -0.073 | 0.129 | 0.244 | 0.247 |
|  | 5mm> | 0.038 | 0.231 | 0.192 | 0.156 | 0.463 | 0.474 |

Differences in the head start position were minimal. When the starting position was at the center of two peaks, the absolute biases between measurers and the IAB method were 0.069 and 0.046 mm, with standard deviations of 0.379 and 0.390 mm, respectively, for the entire length range (Supplementary Table 4). When the starting position was at the upper peak, the absolute biases were 0.101 and 0.052 mm, with standard deviations of 0.369 and 0.380 mm, respectively, for the entire length range (Supplementary Table 4). The absolute bias was smaller in the former case, whereas the standard deviation was larger.

The CCCs were ≥0.95 between measurers in both the ≤5.0- and >5.0-mm ranges (Table 5). For manual measurements vs. the IAB method, the CCCs were ≥0.95 in the ≤5.0-mm range but dropped to <0.90 in the >5.0-mm range (Table 5).

**Table 5.** Concordance correlation coefficients for all samples and for each species. “IAB” denotes the image analysis–based method.

| Body length | A vs B | A vs IAB | B vs IAB |
| --- | --- | --- | --- |
| 5 mm ≤ | 0.986 | 0.961 | 0.957 |
| 5 mm $>$ | 0.982 | 0.861 | 0.862 |

### 3.3 Characteristics of fish showing large length differences between manual measurements and the IAB method

The tip of the head was misdetected in six fish (Figure 7b), and the peak of the head mask was misdetected in three fish (Figure 7c). Moreover, in some larger fish, the masks and curves extended beyond the intended measurement position at the notochord end, reaching the tip of the caudal fin (Figure 7d).

**Figure 7.**
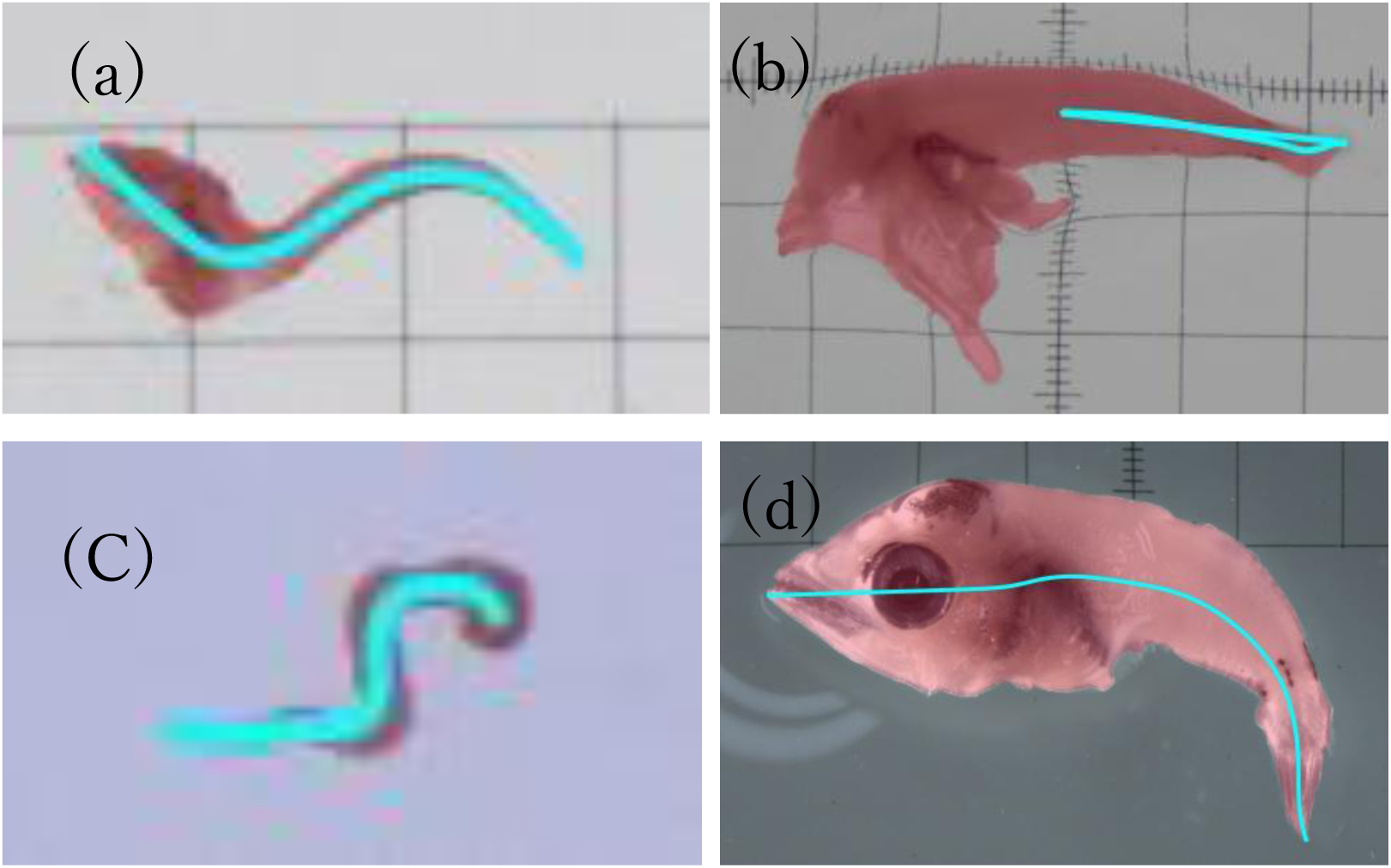
Examples of correct and incorrect curve estimation: (a) correct estimation, (b) misidentification of the tip of the head, (c) failure to detect the head peak, and (d) overmasking of the caudal fin.

## 4. Discussion

This study demonstrated that the curved body length of fixed tuna larvae can be estimated from microscope images with substantial accuracy using image analysis, even without custom training of a deep learning model. However, some fish showed large differences due to errors in the analysis process or changes in body shape during growth. Combining the IAB method with manual remeasurement of fish with large differences, when necessary, enables more efficient length measurements.

### 4.1 Fish region detection

No setting achieved zero values for both under detection and overdetection. Although underdetection could be eliminated, some overdetection remained. Overdetection occurred because of water reflections, debris, plankton, and double detections. Since debris and plankton are often smaller than the minimum fish length (1.5 mm, Supplementary Table 5), applying a length threshold can eliminate some of them (Supplementary Table 6). In addition, these overdetections may be reduced by minimizing the presence of debris and plankton when placing fish on the glass slides. In cases of double detection, an alert for overlapping masks can facilitate subsequent manual remeasurement, although confidence scores or mask area can also be used to select the correct mask (Supplementary Table 7). False detections caused by light reflections can be eliminated by applying a threshold to the proportion of white pixels in the mask, which is typically dominated by these pixels. Changing the box prompt value from 0.30 to 0.35 also reduced the number of overdetections from 49 to 30; however, it resulted in one underdetection. Undetected fish also require searching all images for the missing mask and manual remeasurement. By contrast, most overdetections can be removed or flagged later, although some cannot be processed automatically and must be found manually. The appropriate threshold may depend on the trade-off between manually finding overdetected masks that cannot be processed automatically and manually finding undetected fish, or on the study objective, such as whether all fish or their size composition (i.e., not all, but part of them) is required.

In this study, GroundedSAM detected most tuna larvae without training. However, preparing original data and training a segmentation model is likely necessary to achieve higher detection accuracy. Although the number of images required for training depends on the task and it is difficult to define a standard number for segmentation training, several instance segmentation studies on aquatic organisms have used several thousand individuals or more (Costa et al., 2022; Iwahara et al., 2025; Wang et al., 2025). Because preparing training data for several thousand individuals is labor-intensive, practical implementation will depend on a trade-off between accuracy and the cost of creating training data. However, it will probably be possible to create training data more efficiently by manually refining the automatic masks generated by GroundedSAM.

### 4.2 Curved length estimation

The bias in the differences between manual measurements and the IAB method was equivalent to the bias between measurers in the ≤5.0-mm range. By contrast, the standard deviation of the differences between manual measurements and the IAB method was higher than that between measurers. Although the IAB method accurately estimated the length of most tuna larvae, particularly in the ≤5.0-mm range, the errors in the curved length estimation process were likely one reason for the higher standard deviation. According to the strength-of-agreement criteria for the CCC, the IAB method demonstrated substantial agreement in the ≤5.0-mm range (McBride and others, 2005). However, determining an acceptable margin of difference is complex and can vary depending on the purpose for using length data and how it is used (Bunch et al., 2013; Elstner et al., 2025). For cases requiring higher accuracy than the IAB method can provide, manually checking and remeasuring these fish can maintain accuracy while improving measurement efficiency. In addition to accuracy-related limitations, the IAB method cannot support fish with partially collapsed bodies, looped bodies, or three-dimensional bends. Therefore, manual measurement is required for these types of fish.

A bimodal distribution of differences between manual measurements and the IAB method was observed in fish sizing >5 mm: some fish were estimated accurately, whereas others showed increasing differences as their size increased. This bimodal distribution is likely to be linked to the extension of the caudal fin during growth. In smaller fish, the caudal fin is barely formed at the notochord end, which is the terminal point of the length measurement, and it extends as the fish grows. However, even in larger fish, the caudal fin remained unclear in images, which probably resulted in two patterns: one in which the caudal fin was masked by GroundedSAM (Supplementary Figure 6a) and another in which it was not masked (Supplementary Figure 6b). Since the proper endpoint is the notochord end, accurately detecting it in larger fish becomes necessary. To address this issue, training a model with custom data may be necessary, such as a segmentation model to avoid masking regions extending to the caudal fin or a keypoint model to identify the position of the notochord end and use it as the endpoint. However, 72.5% of the fish sampled using ring nets in Japanese coastal waters from 2017 to 2024 were smaller than 5.0 mm (Supplementary Table 5). This indicates that our current method can still improve the efficiency of this measurement task to some extent. For example, when visualizing the estimated curves, using a color change to highlight fish sizing >5 mm could facilitate manual remeasurement and improve overall measurement efficiency.

The difference in the measurement starting point did not lead to a notable difference when either the peak center or the upper peak was selected. However, because our current method cannot identify the upper jaw, using the upper peak as the starting point requires placing the upper jaw toward the top of the image. This positioning task requires meticulous effort, so the decision on which starting point to use depends on the trade-off between adhering to the correct anatomical measurement position and reducing the workload.

### 4.3 Future applications

Full automation of this method requires automatic scale detection. We used previously captured images with a large number of samples to achieve ideal sampling for evaluation. However, because these images were intended for manual length measurement using tools such as ImageJ, the scales were graduated or grid-marked glass slides. These scales were not well suited for automatic detection, so we manually extracted the scale information in this study. In future, printing or placing colored, fixed-size scales (e.g., Shibata et al., 2024) on glass slides will enable automatic scale detection and fully automated measurements.

This method has the potential to be widely applied to the larvae of other species. Although this study evaluated only five species, no species-specific training was required. The method can be used to measure notochord length and standard length in larvae at stages when the caudal fin has not yet extended. However, the applicable body-length range may vary among species, because the body length at which the caudal fin begins to extend probably differs by species. Furthermore, although this study found little difference in the starting point when multiple peaks appeared in the head mask—whether using the center peak or the upper peak—this method should be applied with caution to species in which the upper and lower jaws differ markedly in length (e.g., family Holocentridae; Muneo, 2014).

## 5. Conclusion

This study demonstrated that the curved body length of most tuna larvae acquired in microscope-captured images can be estimated with practical accuracy, even without prior training. However, achieving detection with both underdetection and overdetection reduced to zero was difficult. In addition, training a segmentation or keypoint model with custom data would be necessary to further improve accuracy, particularly for larger fish with a caudal fin. Nevertheless, semi-automated measurement, combining the IAB method with manual correction when needed, can enhance the efficiency of length measurement tasks.

## Acknowledgments

We thank the captains and crews of the R/V Kaiyo-maru, Shunyo-maru, and Yoko-maru for larval sampling, and the scientists at the Fisheries Resources Institute for their cooperation with the sampling. We would also like to thank Mr. Hirohito Miyashita for providing high-quality annotated data.

## Data availability

The authors do not have permission to share data.

