## Supplementary materials for "Estimation of the curved body length of tuna larvae from microscope images using a zero-shot model and image processing techniques"

Supplementary material captions

Supplementary Table 1. Number of fish used per 0.5-mm-size class for each species. Common and scientific names of the five species: Pacific bluefin tuna *Thunnus orientalis*, yellowfin tuna *Thunnus albacares*, skipjack tuna *Katsuwonus pelamis*, bullet tuna *Auxis rochei*, and frigate tuna *Auxis thazard*.

Supplementary Table 2. Shapiro–Wilk test results assessing data normality. All data were significantly non-normal. *$p$-value < 0.05.

Supplementary Table 3. Bias and standard deviation of the relative differences between measurers A and B, and between each measurer and the IAB method (IAB) under the three aggregation conditions (the entire length range, ≤5 mm, and >5 mm).

Supplementary Table 4. Bias and standard deviation of the differences between measurers A and B, and between each measurer and the IAB method (IAB) under the three aggregation conditions (the entire length range, ≤5 mm, and >5 mm). The IAB method included two approaches: one in which the starting point of the head was the average of the two peaks (Center) and another in which it was the upper peak (Upper).

Supplementary Table 5. Body length composition of the full dataset (21,836 individuals of five tuna species sampled using ring nets in Japanese coastal waters, 2017–2024). Percentages represent the proportion of each 0.5-mm body-length bin relative to the total sample (100%).

Supplementary Table 6. Lengths of overdetected debris, sorted in ascending order.

Supplementary Table 7. Area and score for double-detected cases. * indicates cases where the correct mask can be selected by choosing the larger score or the smaller area. “Total” indicates the number of such cases.

Supplementary Figure 1. Examples of the images and scale markings used in this study.

Supplementary Figure 2. Images of the notochord length and standard length. The notochord and standard body lengths refer to the same anatomical region, whose name varies with the growth stage.

Supplementary Figure 3. Examples of fish positioning. (a) Correct positioning: the fish is oriented with the upper jaw at the top. (b) Incorrect positioning: the fish is oriented with the upper jaw at the bottom.

Supplementary Figure 4. Scatter plots of body length versus relative differences: (a) measurers A and B, (b) measurer A vs. the IAB method, and (c) measurer B vs. the IAB method. Common and scientific names of the five species: Pacific bluefin tuna *Thunnus orientalis*, yellowfin tuna *Thunnus albacares*, skipjack tuna *Katsuwonus pelamis*, bullet tuna *Auxis rochei*, and frigate tuna *Auxis thazard*.

Supplementary Figure 5. Scatter plot showing body length and measurement differences when the tip of the upper jaw was used as the tip of the head: (a1–a3) relative differences and (b1–b3) absolute differences. (1) Measurers A and B, (2) measurer A vs. the IAB method, and (3) measurer B vs. the IAB method. Common and scientific names of the five species: Pacific bluefin tuna *Thunnus orientalis*, yellowfin tuna *Thunnus albacares*, skipjack tuna *Katsuwonus pelamis*, bullet tuna *Auxis rochei*, and frigate tuna *Auxis thazard*.

Supplementary Figure 6. Examples of masks and curves for large fish (>5 mm). Red regions indicate GroundedSAM masks, and blue lines indicate the estimated curves. (a) With the caudal fin clearly visible, GroundedSAM generated a mask extending to the fin, causing the curve and the measurement position to reach the fin tip and resulting in an overestimation of length. A blue point indicates the notochord end, the correct endpoint. (b) When the caudal fin was indistinct, the mask did not extend to it, and the curve was correctly generated up to the notochord end.

**Supplementary table 1**

| Bin width (mm) | Number | | | | |
|---|---|---|---|---|---|
| | Pacific bluefin tuna | Yellow tuna | Skipjack tuna | Bullet tuna | Frigate tuna |
| 1.5-2.0 | 20 | | | | |
| 2.0-2.5 | 20 | 20 | 20 | 20 | |
| 2.5-3.0 | 20 | 20 | 20 | 20 | |
| 3.0-3.5 | 20 | 20 | 20 | 20 | 20 |
| 3.5-4.0 | 20 | 20 | 20 | 20 | 20 |
| 4.0-4.5 | 20 | 20 | 20 | 20 | 20 |
| 4.5-5.0 | 20 | 20 | 20 | 20 | 20 |
| 5.0-5.5 | 20 | 20 | 20 | 20 | 20 |
| 5.5-6.0 | 20 | 20 | 20 | 20 | 20 |
| 6.0-6.5 | 20 | 20 | 20 | 20 | 20 |
| 6.5-7.0 | 20 | 20 | | 20 | |
| 7.0-7.5 | 20 | 20 | | | |
| 7.5-8.0 | 20 | 20 | | | |
| 8.0-8.5 | 20 | | | | |

**Supplementary table 2**

| | | Relative bias | | | | Absolute bias | | | |
|---|---|---|---|---|---|---|---|---|---|
| | | A vs IAB | | B vs IAB | | A vs IAB | | B vs IAB | |
| Shapiro–Wilk test | W | 0.943 | | 0.948 | | 0.925 | | 0.939 | |
| | P value | 6.65E-16 | * | 3.42E-15 | * | 2.20E-16 | * | 2.20E-16 | * |

**Supplementary table 3**

| Species | Body length range | Relative bias (%) | | | Standard deviation of relative difference (%) | | |
|---|---|---|---|---|---|---|---|
| | | A vs B | A vs IAB | B vs IAB | A vs B | A vs IAB | B vs IAB |
| All | All | 1.308 | 1.203 | -0.019 | 3.375 | 7.442 | 7.700 |
| | 5mm<= | 1.837 | -0.661 | -2.365 | 3.855 | 7.387 | 7.453 |
| | 5mm> | 0.658 | 3.491 | 2.859 | 2.527 | 6.855 | 6.996 |

**Supplementary table 4**

| Head point | Body length range | Absolute bias (mm) | | | Standard deviation of absolute difference (mm) | | | Relative bias (%) | | | Standard deviation of relative difference (%) | | |
|---|---|---|---|---|---|---|---|---|---|---|---|---|---|
| | | A vs B | A vs IAB | B vs IAB | A vs B | A vs IAB | B vs IAB | A vs B | A vs IAB | B vs IAB | A vs B | A vs IAB | B vs IAB |
| Center | All | 0.049 | 0.096 | 0.046 | 0.142 | 0.379 | 0.390 | 1.308 | 1.203 | -0.019 | 3.375 | 7.442 | 7.700 |
| | 5mm<= | 0.059 | -0.014 | -0.073 | 0.129 | 0.244 | 0.247 | 1.837 | -0.661 | -2.365 | 3.855 | 7.387 | 7.453 |
| | 5mm> | 0.038 | 0.231 | 0.192 | 0.156 | 0.463 | 0.474 | 0.658 | 3.491 | 2.859 | 2.527 | 6.855 | 6.996 |
| Upper | All | 0.049 | 0.101 | 0.052 | 0.142 | 0.369 | 0.380 | 1.308 | 1.357 | 0.131 | 3.375 | 7.166 | 7.423 |
| | 5mm<= | 0.059 | -0.007 | -0.065 | 0.129 | 0.232 | 0.238 | 1.837 | -0.422 | -2.131 | 3.855 | 7.050 | 7.114 |
| | 5mm> | 0.038 | 0.234 | 0.196 | 0.156 | 0.454 | 0.464 | 0.658 | 3.539 | 2.906 | 2.527 | 6.696 | 6.835 |

**Supplementary table 5**

| Length (mm) | Composition (%) |
|---|---|
| 1.5-2.0 | 0.44 |
| 2.0-2.5 | 5.23 |
| 2.5-3.0 | 8.42 |
| 3.0-3.5 | 10.07 |
| 3.5-4.0 | 11.76 |
| 4.0-4.5 | 17.84 |
| 4.5-5.0 | 18.73 |
| 5.0-5.5 | 13.95 |
| 5.5-6.0 | 7.95 |
| 6.0-6.5 | 2.89 |
| 6.5-7.0 | 1.27 |
| 7.0-7.5 | 0.67 |
| 7.5-8.0 | 0.42 |
| 8.0-8.5 | 0.22 |
| 8.5-9.0 | 0.09 |
| 9.0-9.5 | 0.05 |
| 9.5-10.0 | 0.00 |

**Supplementary table 6**

| Number | Length (mm) |
|---|---|
| 1 | 0.35 |
| 2 | 0.58 |
| 3 | 0.62 |
| 4 | 0.66 |
| 5 | 0.67 |
| 6 | 0.68 |
| 7 | 0.69 |
| 8 | 0.69 |
| 9 | 0.70 |
| 10 | 0.79 |
| 11 | 0.92 |
| 12 | 0.96 |
| 13 | 1.00 |
| 14 | 1.21 |
| 15 | 1.26 |
| 16 | 1.35 |
| 17 | 1.38 |
| 18 | 1.43 |
| 19 | 1.81 |
| 20 | 2.25 |

**Supplementary table 7**

| Score | | | | Area（pixels） | | | |
|---|---|---|---|---|---|---|---|
| Correct | | Incorrect detection | | Correct | | Incorrect detection | |
| 0.36 | | 0.41 | - | 140,854 | * | 170,611 | - |
| 0.43 | * | 0.37 | - | 193,501 | * | 199,150 | - |
| 0.43 | * | 0.36 | - | 24,781 | * | 25,288 | - |
| 0.3 | | 0.49 | - | 28,236 | | 28,171 | - |
| 0.39 | * | 0.38 | - | 290,794 | * | 291,756 | - |
| 0.45 | * | 0.37 | - | 242,975 | * | 245,685 | - |
| 0.3 | | 0.5 | - | 344,682 | * | 346,272 | - |
| 0.31 | | 0.31 | 0.45 | 28,084 | | 14,155 | 10,837 |
| 0.36 | | 0.44 | - | 131,015 | * | 132,828 | - |
| 0.37 | | 0.39 | - | 122,095 | * | 122,264 | - |
| 0.43 | * | 0.43 | - | 50,019 | * | 51,024 | - |
| 0.32 | | 0.45 | - | 766,221 | * | 807,679 | - |
| 0.44 | * | 0.38 | - | 291,069 | * | 292,412 | - |
| 0.42 | * | 0.34 | - | 301,544 | * | 302,863 | - |
| 0.5 | * | 0.31 | - | 459,678 | * | 461,560 | - |
| 0.35 | | 0.5 | - | 186,413 | * | 186,736 | - |
| 0.44 | * | 0.36 | - | 356,033 | * | 361,093 | - |
| 0.44 | * | 0.39 | - | 1,425,519 | * | 1,430,066 | - |
| Total | 10 | | | Total | 18 | | |

8

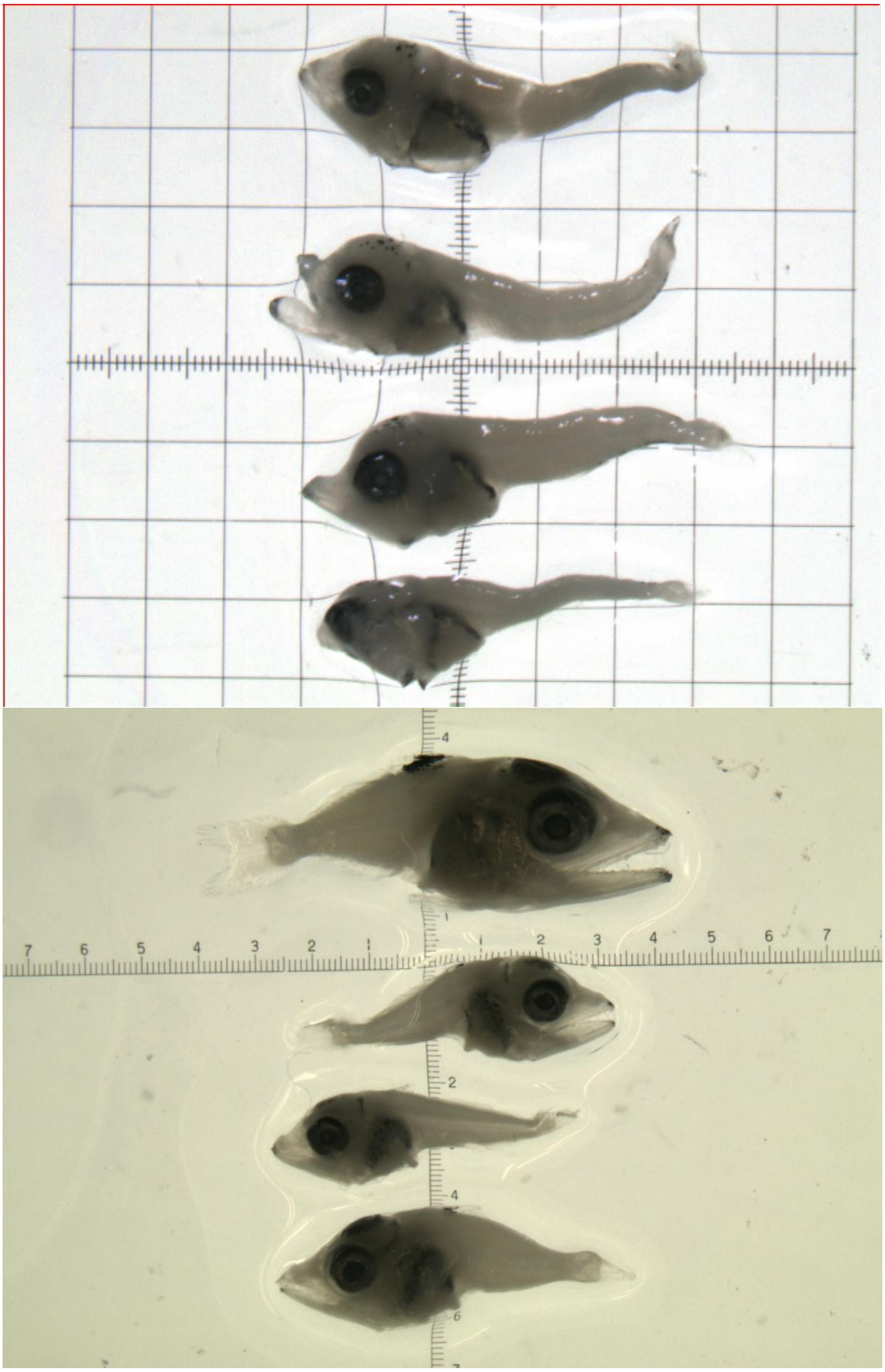

**Supplementary figure 1**

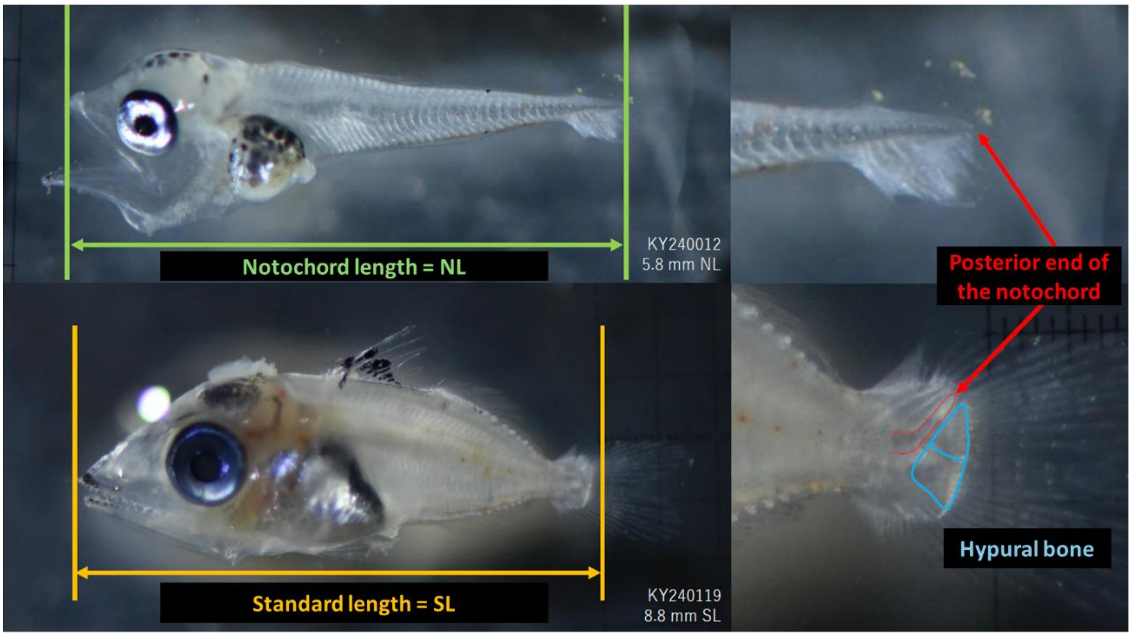


**Supplementary figure 2**

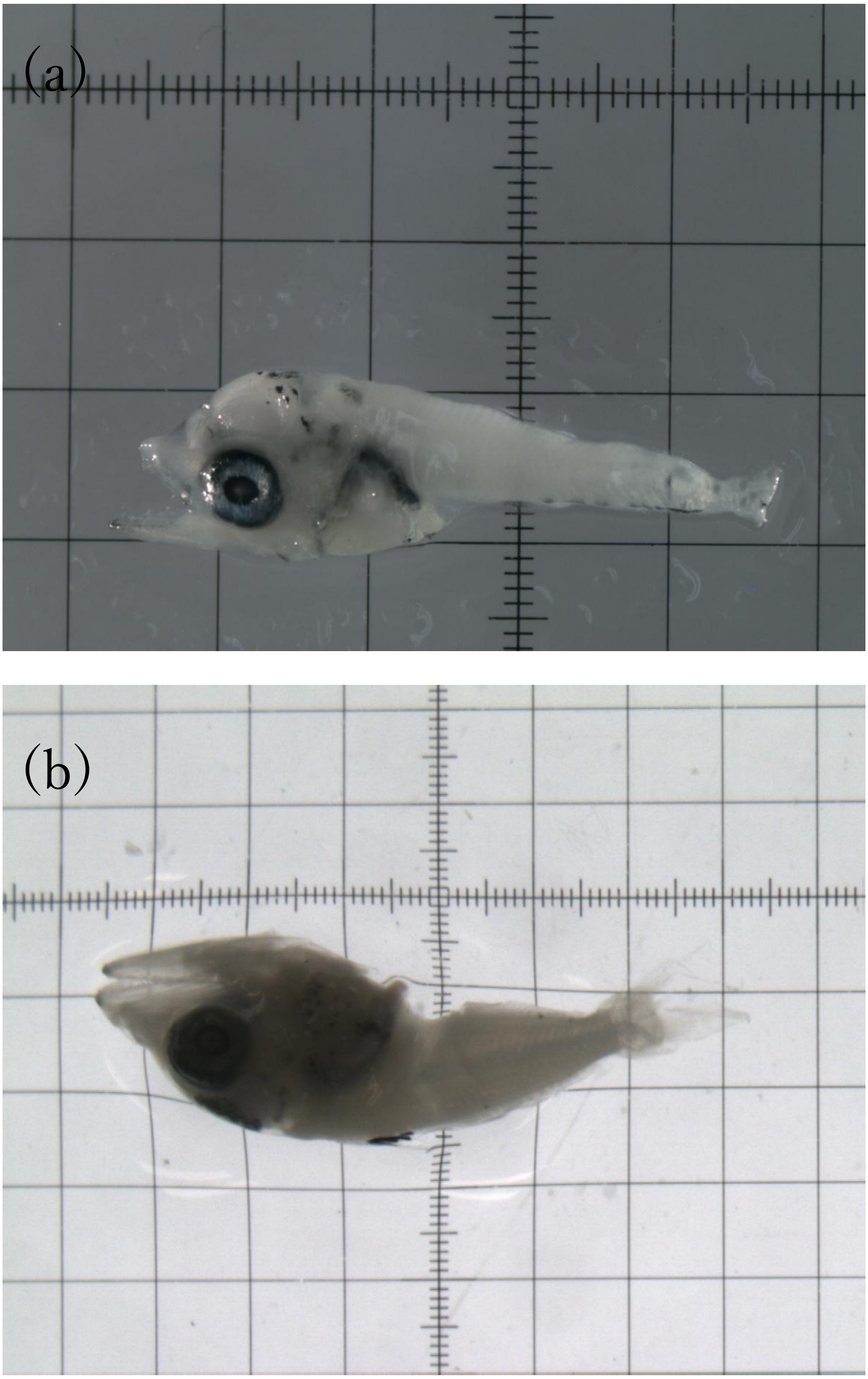


**Supplementary figure 3**

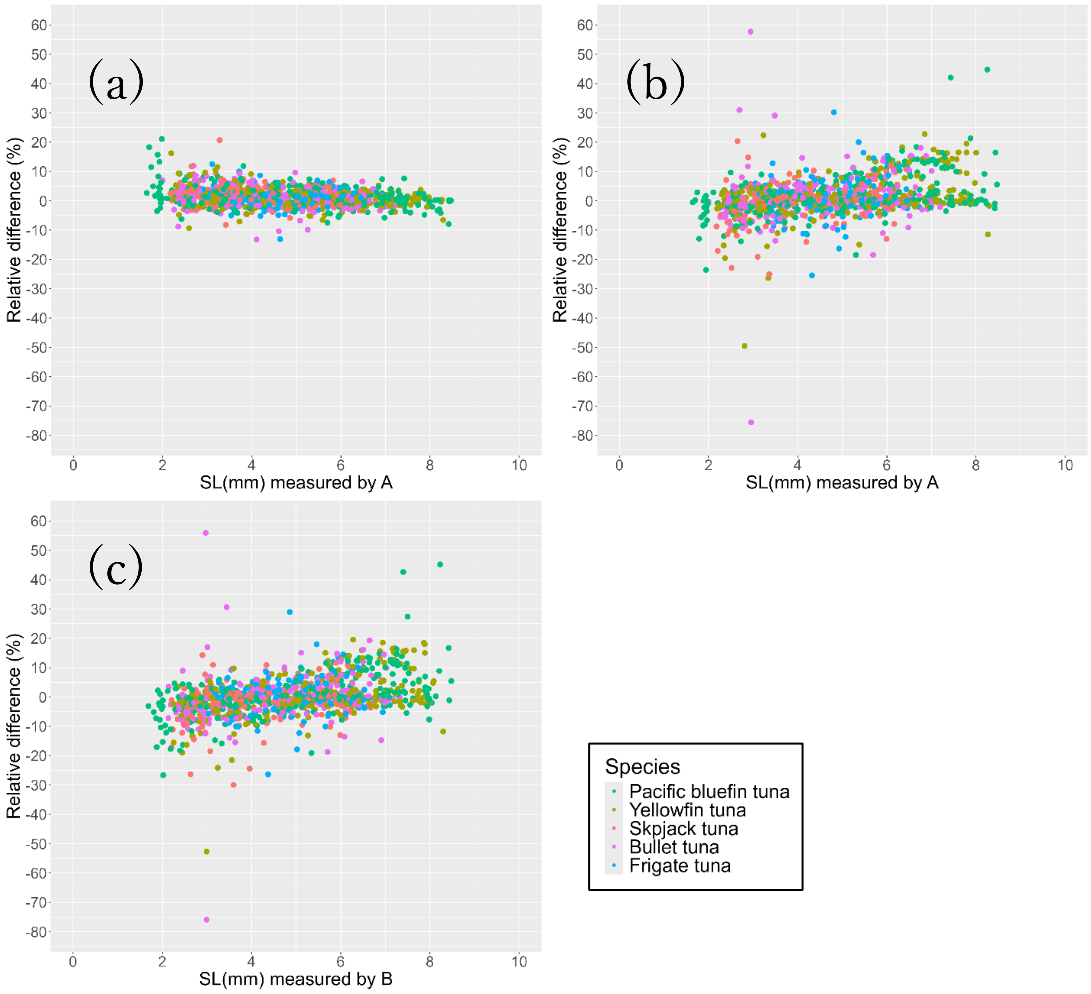


**Supplementary figure 4**

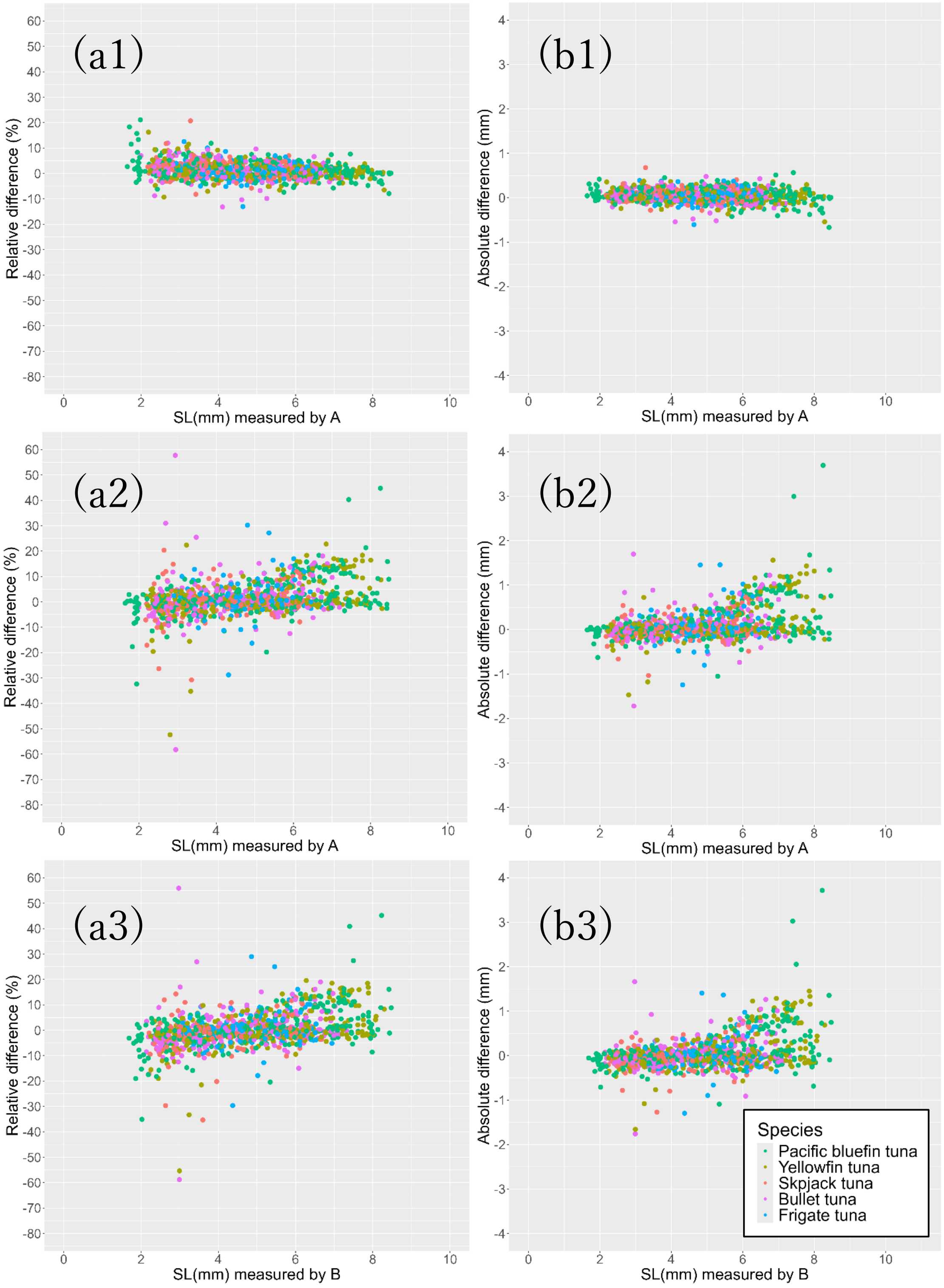


20

21 **Supplementary figure 5**

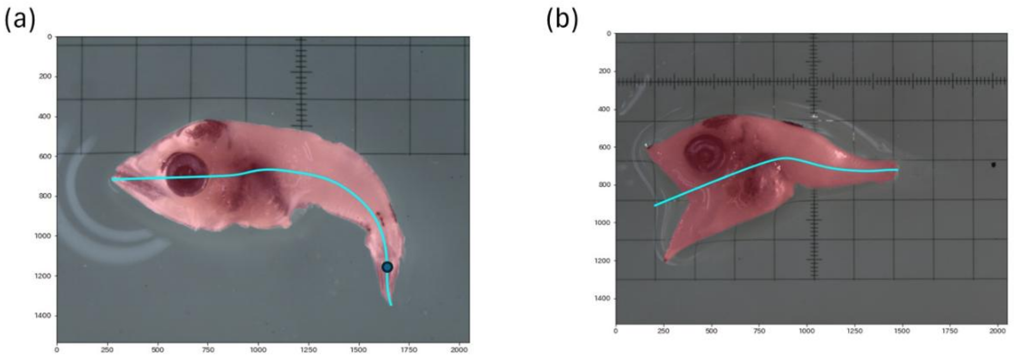


**Supplementary figure 6**
